# The interplay between habitat fragmentation and life traits affects specialization in lichen symbioses

**DOI:** 10.1101/2022.03.30.485788

**Authors:** Alejandro Berlinches de Gea, Miguel Verdú, Mar Villar-dePablo, Sergio Pérez-Ortega

## Abstract

Interactions between organisms are determined by species traits and differ in specialization, from generalist to highly specialized relationships. Although we expect that the effect of habitat loss and fragmentation on the abundance and survival of species depends on their degree of specialization, few studies have deepened into the interplay between interaction specialization and habitat loss.
Here, we investigate the combined effect of habitat fragmentation and life traits (growth type and reproductive mode) on the specialization of lichen-forming fungi (mycobionts) towards its photosynthetic partners (photobionts) in lichen symbioses.
We studied mycobiont specialization in epiphytic lichen communities present in ten fragments of *Quercus rotundifolia* trees embedded in an agricultural matrix. Both mycobionts and photobionts were identified through DNA sequencing and mycobiont specialization was measured through interaction parameters calculating the relative number of interactions (normalized degree; ND) and the specialization of each species based on its discrimination from a random selection of partners (d’). Phylogenetic generalized linear mixed models were used to analyze the effect of patch size as well as the life traits growth type (crustose, foliose, fruticose) and reproduction mode (sexual vs asexual) on mycobiont specialization.
Both mycobiont and photobiont richness along the patch size gradient followed a hump-back pattern, which was more pronounced in photobionts. Mycobionts forming crustose biotypes established the largest number of interactions. Mycobiont specialization (d’) was larger for fruticose and foliose biotypes and for species with vegetative reproduction. Along the gradient of fragment size, the relative number of interactions decreased and the specialization of mycobionts with vegetative reproduction increased.
*Synthesis*: The analysis of mycobiont specialization towards their photobionts in communities of epiphytic lichens in a fragmented Mediterranean forest revealed that the interaction between species life traits and habitat loss on specialization cannot be neglected. The results also pointed to the ability of some species to modulate their interactions to certain extent, suggesting that species might have a greater resilience to abiotic changes than expected, presumably creating extinction debt or even avoiding extinction processes to some extent.

**Resumen:**

1. Las interacciones entre organismos están determinadas por los rasgos funcionales de las especies y difieren en su grado de especialización, desde interacciones generalistas hasta las altamente especializadas. Aunque se espera que el efecto de la pérdida y fragmentación del hábitat sobre la abundancia y la supervivencia de las especies dependa de su grado de especialización, pocos estudios han profundizado en la interacción entre la especialización de las interacciones y la pérdida de hábitat.
2. En este artículo investigamos el efecto combinado de la fragmentación del hábitat y los rasgos funcionales de las especies (tipo de crecimiento y modo reproductivo) en la especialización de los hongos liquenizados (micobiontes) hacia sus socios fotosintéticos (fotobiontes) en las simbiosis de líquenes.
3. Se estudió la especialización de micobiontes en comunidades de líquenes epífitos presentes en diez fragmentos de árboles de *Quercus rotundifolia* incrustados en una matriz agrícola. Tanto los micobiontes como los fotobiontes fueron identificados mediante la secuenciación del ADN y la especialización de los micobiontes se midió a través de parámetros de interacción calculando el número relativo de interacciones (grado normalizado; ND) y la especialización de cada especie en función de su discriminación de una selección aleatoria de simbiontes (d’). Se utilizaron modelos lineales mixtos generalizados filogenéticos para analizar el efecto del tamaño del parche, así como los rasgos de tipo de crecimiento (crustáceo, folioso, frutal) y el modo de reproducción (sexual frente a asexual) sobre la especialización de los micobiontes.
4. Tanto la riqueza de micobiontes como de fotobiontes a lo largo del gradiente de tamaño de los parches siguió un patrón de U inversa, que fue más pronunciado en los fotobiontes. Los micobiontes que forman biotipos crustáceos establecieron el mayor número de interacciones. La especialización de los micobiontes (d’) fue mayor para los biotipos fruticulosos y foliosos y para las especies con reproducción vegetativa. A lo largo del gradiente del tamaño del fragmento, el número relativo de interacciones disminuyó y la especialización de los micobiontes con reproducción vegetativa aumentó.
5. Síntesis: El análisis de la especialización de los micobiontes hacia sus fotobiontes en comunidades de líquenes epífitos en un bosque mediterráneo fragmentado reveló que no se puede descartar la interacción entre los rasgos de vida de las especies y la pérdida de hábitat sobre la especialización. Los resultados también apuntaron a la capacidad de algunas especies para modular sus interacciones hasta cierto punto, sugiriendo que las especies podrían tener una resiliencia a los cambios abióticos mayor de lo esperado, presumiblemente creando una deuda de extinción o incluso evitando los procesos de extinción hasta cierto punto. Palabras clave: Bosque mediterráneo, epífitos, fotobiontes, hongos liquenizados, selectividad.

## Introduction

Habitat loss due to anthropogenic land use represents one of the main drivers of biodiversity loss under the current scenario of global change (Haddad et al., 2015). A common process following habitat loss is habitat fragmentation in which natural homogeneous habitat is transformed into smaller fragments embedded in a matrix different from the original one (Wilcove et al., 1986). Consequently, fragmentation leads to a reduction in the amount of available habitat for species as well as its connectivity, and to changes of abiotic and biotic conditions in the remaining habitat (Fahrig, 2003). This process usually entails the decrease of population sizes, increasing the risk of species extinctions, and subsequent reduction of ecosystem structure, functioning and services (Isbell et al., 2011; Millennium ecosystem assessment 2005).

Changes derived from habitat fragmentation affect species unevenly depending on life strategies, population sizes and reproductive and physiological traits (Bovo et al., 2018; Farneda et al., 2015; Henle et al., 2004). Ecological specialization towards abiotic and biotic conditions is a highly relevant trait that may determine species fate during habitat fragmentation, being specialists more prone to suffer fragmentation effects (Devictor et al., 2008; Nordén et al., 2013).

As species interact with other species in natural habitats, fragmentation and habitat loss jeopardize species interactions (Tylianakis et al., 2008). Whereas the loss of taxonomic diversity has received large attention during the last decades, knowledge on the effect of habitat loss on interactions is still scarce (Gonzalez et al., 2011). The loss of interactions often precedes the loss of species diversity in ecosystems and it is seen as an early signal of ecosystem decay (Aizen et al., 2012; Valiente-Banuet et al., 2015). Not all kinds of interactions are equally affected by fragmentation, which has been shown to produce more negative effects on mutualisms than on antagonisms (Magrach et al., 2014). Most of the knowledge we have on the effect of fragmentation on mutualisms comes from non-intimate interactions (e.g., seed dispersal, plant pollination) whereas intimate mutualisms have been largely ignored.

Independently of the interaction type, species show variability regarding the degree of specialization towards their partners, which ranges from generalists, interacting with many partners, to specialists, which interact with a few or even a single partner (Solé and Montoya 2001; Thompson, 1988). These strategies involve different evolutionary and ecological advantages and constraints (Dennis et al., 2011; Ollerton, 2006). Furthermore, they have been highly relevant to predicting the loss of interactions and species in gradients of habitat availability (Aizen et al., 2012). In addition, specialization is correlated with morphological and reproductive traits (Maglianesi et al., 2014; Otálora et al., 2013; Reif et al., 2016; Santamaría and Rodríguez-Gironés, 2007) as well as with species’ evolutionary history (Webb et al., 2010). However, it is largely ignored whether species might modulate their specialization under changing environmental conditions in order to survive. Overall specialization tends to decrease together with habitat availability (Hadley et al., 2018; Jauker et al., 2019). This decrease may be due to the loss of specialists (Aizen et al., 2012; Bagchi et al., 2018) or due to the variation of specialization within species, which may show certain plasticity to cope with new habitat conditions.

Lichens, the symbiotic phenotype of lichen-forming fungi (hereafter, the mycobiont) associated with, at least, one photosynthetic partner (the photobiont) a cyanobacteria and/or a green alga (Grube and Hawksworth 2007) are paradigmatic examples of intimate mutualisms. Due to their high sensibility to subtle changes in abiotic conditions in the environment, they are renowned bioindicators of habitat conditions (Marmor et al., 2011; McCune 2000; Nascimbene and Marini 2015; Rivas Plata et al., 2008), including changes in land management and habitat fragmentation (Aragón et al., 2010; Brunialti et al., 2013; Matos et al., 2017; Svoboda et al., 2010).

Some functional traits of lichens, mostly related to growth types and reproductive modes, are commonly used in ecological and indicator studies as a response to many environmental or anthropogenic changes (Ellis et al., 2021; Giordani et al., 2012; Hurtado et al., 2020; Koch et al., 2019; Nelson et al., 2015). Another functional trait commonly used is the photobiont type, often categorized as highly broad taxonomic categories (e.g. Giordani et al., 2012). The last two decades have seen great advances in the understanding of photobiont diversity and the range of fungal-algal interactions in lichen symbioses (Muggia et al., 2020), especially to know the breadth of compatible photobionts at different systematic levels (e.g. (Dal Grande et al., 2012; Fernández-Mendoza et al., 2011; Leavitt et al., 2015; Thüs et al., 2011)). During this time, it has become clear that mycobiont specialization towards the photobiont varies among species and it is likely a species-specific trait (Magain et al., 2016; Pérez-Ortega et al., 2012). However, the knowledge about the influence of certain functional traits on photobiont specialization is scarce.

Lichens show a wide range of reproductive strategies (Tripp and Lendemer, 2018), but they can be simply divided between asexual versus sexual reproduction. Most common asexual reproductive strategies involve vegetative clonal propagules (soredia, isidia…) in which both bionts are dispersed together ensuring the availability of compatible photobionts for the fungi at the expense of suppressing genetic variability in both bionts (Buschbom and Mueller 2005; Yahr et al., 2006). On the contrary, the dispersal of the mycobiont via ascospores implies the propagule to find a compatible photobiont after its establishment on the new substrate (Fernández-Mendoza et al., 2011; Yahr et al., 2006). The scarce studies in which specialization towards the photobiont has been explored within a reproductive framework offer contradictory results (Otálora et al., 2013; Wornik and Grube, 2010), likely due to the possibility of photobiont switches after the vegetative propagule is settled (Nelsen and Gargas, 2008)

Lichen thalli also show a number of growth forms named biotypes (Honegger, 2001), which have numerous implications for associated photobionts since different architectures and topologies imply the existence of different microniches (Hawksworth and Grube, 2020). The most common biotypes, which are the focus of this study, are: 1) crustose, in which the thallus lacks a lower cortex and is attached to the substrate through the medulla, 2) foliose, in which the thallus has a laminar form, partially adhered to the substrate by certain fixation structures, 3) fruticose, in which thallus shows a shrub-like form attached to the substrate usually by a single point. The role of thallus morphologies on photobiont specialization has not been formally tested, although it is known that, for instance, crustose species usually display a high phylogenetic breadth of photobiont partners (Blaha et al., 2006; Guzow-Krzeminska, 2006; Muggia et al., 2014).

In this study, we explore the effect of habitat fragmentation on mycobiont specialization toward their photobionts as well as the effect of growth and reproduction traits on the specialization along the fragmented habitat. We study the richness and specialization of epiphytic lichen-forming fungi and their associated photobionts in habitat fragments of Mediterranean forest of increasing size. We hypothesize that habitat loss reduces both biont richness and specialization. Finally, we explore whether species specialization may be affected by species-specific functional traits such as reproductive mode and growth form.

## Material and Methods

### Study area

The area of study was located in Central Spain (42° 5-8’ N; 3° 36-44’ W; 923-1093 m a.s.l.). The area harbors remnants of forest embedded in an agricultural matrix (cereal crops) (Figure 1). Fragments are either dominated by one of the main dominating tree species in the area (*Quercus rotundifolia, Q. pyrenaica* and *Q. faginea*) or show mixed formations in which *J. oxycedrus* also occasionally occur.

**Figure 1.**
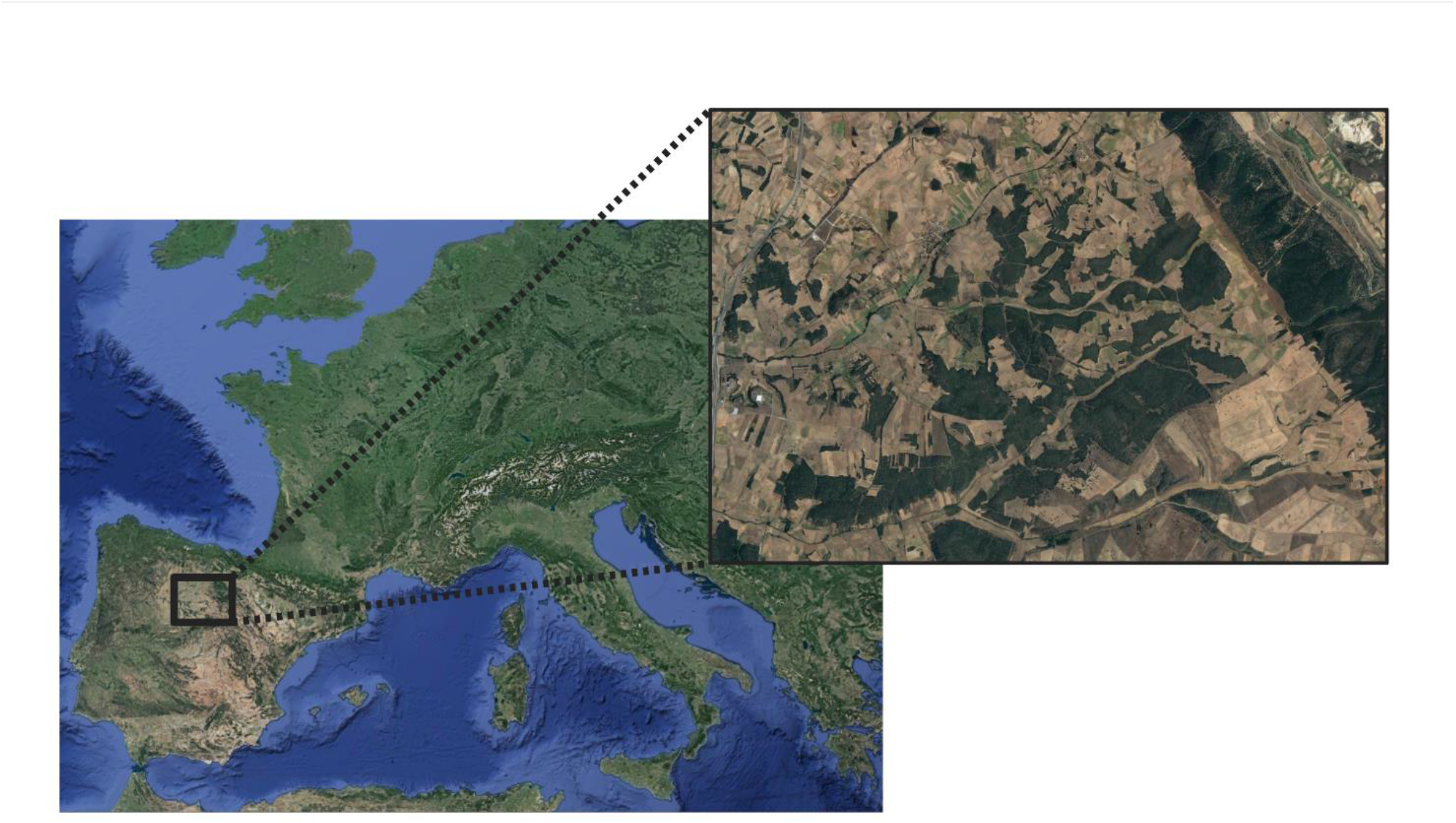
Map detailing the area where the study was carried out. Images obtained from Google Earth (14/12/2015) Europe. (Landsat/Copernicus Data SIO, NOAA, U.S. Navy, NIGA, GEBCO, GeoBasis-DE/BKG (©2009) Map GISrael Instituto Geográfico Nacional; 45°09’57”N 11°06’33”E. https://earth.google.com/web/@45.16589093,11.10934345,1020.81870028a,3304630.75835109d,35y,0h,0t,0r) The second map (smaller one on the right) is also obtained from Google Earth (20/08/2018). Madrigal del Monte. Instituto Geográfico Nacional. 42°07’17”N 3°38’41”W. https://earth.google.com/web/@42.12151496,3.64472552,955.37218271a,9380.0627212d,35y,0h,0t,0r

### Sampling

The selection of the habitat fragments was made as follows: Orthorectified photographs were downloaded from the National Centre for Geographic Information (http://centrodedescargas.cnig.es/CentroDescargas/-index.jsp) corresponding to the year 2017, and by means of the Google Earth Pro software (available at https://www.google.com/earth/download/gep/agree.html) all forest fragments present in the area were identified. Perimeter and area were measured for each fragment. Fragments with highly irregular geometries (low area/perimeter ratio) were discarded to avoid disproportionate edge effects. The remaining fragments were inspected during the fall of 2017 in order to check the tree composition homogeneity. Considering the available diversity of fragment sizes, we chose fragments in which *Quercus rotundifolia* was the dominant species (> 75 %). Finally, a total of 10 fragments were selected in a size gradient ranging from 0.002 ha (a single isolated tree) to 250ha (Table 1).

**Table 1.** Mycobiont and photobiont richness and mean mycobiont specialization in each studied forest patch. A= Area of each forest patch, in hectares; M_d_= Number of mycobiont species; M_e_= Number of exclusive mycobiont species in a forest patch; P_d_= Number of photobiont ASVs; P_e_= Number of exclusive ASV species in a forest patch; d= mean degree; ND: mean normalized degree.

| A | M<sub>d</sub>/M<sub>e</sub> | P<sub>d</sub>/P<sub>e</sub> | d | ND |
| --- | --- | --- | --- | --- |
| 0.002 | 9/1 | 25/3 | 9 | 0.33 |
| 0.1 | 17/1 | 34/8 | 7.64 | 0.22 |
| 0.3 | 22/1 | 42/6 | 9.13 | 0.217 |
| 1.02 | 23/1 | 63/12 | 10.83 | 0.17 |
| 4.23 | 27/1 | 51/13 | 8.37 | 0.17 |
| 12.21 | 28/1 | 75/20 | 9.54 | 0.13 |
| 16.13 | 23/0 | 57/8 | 11.87 | 0.21 |
| 35.85 | 23/1 | 52/9 | 8.39 | 0.16 |
| 48.01 | 27/0 | 61/12 | 8.89 | 0.15 |
| 250 | 20/2 | 46/7 | 2.9 | 0.06 |

In each fragment, epiphytic lichen communities were surveyed at the geographic center of the fragments. Sampling was carried out according to Asta et al., (2002). In summary, we sampled the lichen community in a total of ten tree trunks (diameter ≥ 12 cm) in both north and south orientation, using a 10×50 cm grid divided into five 10×10 cm squares and annotated all species present and their abundance which was measured as the number of squares where the species occurs. Once the inventory was completed, we attempted to collect a total of 10 individuals of all the species identified in each fragment. This search lasted up to a maximum of 4 hours per plot (two people). Samples were dried at room temperature and stored until further processing.

### Molecular methods

#### Mycobiont barcoding

Field identifications of mycobionts were subsequently checked using currently used methods in lichen taxonomy and available literature. Further, we sequenced the nrITS region, the universal fungal barcode for fungi (Schoch et al., 2012) to corroborate identifications in each of the identified morphospecies. Briefly, DNA extractions were performed using cationic exchange resin Chelex 100 (BioRad, Madrid) following Ferencova et al., (2017). PCRs were performed using the primers ITS1F and ITS4-Kyo2 (Gardes and Bruns 1993; Toju et al., 2012). Reactions included 6.5 μl of MyTaq Red Mix (BioLine, UK), 2 μl of diluted (1:10) DNA template, 0.5 μl (10uM) of each primer and 5.5 μl of sterile distilled H_2_O. PCR conditions were set as follows: a denaturation step of 94°C for 5 min followed by 35 cycles of 95°C for 30 s, 57°C for 30 s, and 72 °C for 1 min ending with an elongation of 72°C for 7 min. Amplicons were sequenced in Macrogen (Madrid, Spain) using the same primers as in the amplification reaction.

#### Mycobiont phylogeny

We constructed the phylogenetic tree depicting the evolutionary relationships of mycobionts to account for statistical non-independence derived from common ancestry in the models. DNA extractions were either using extractions obtained in the previous section or using E.Z.N.A.® Forensic Kit (Omega Bio-Tek, Norcross, Georgia, USA) according to the protocol defined by the company. To determine phylogenetic affinities, we used DNA three nuclear genomic regions, *i*.*e*. the ribosomal internal transcribed spacer (nrITS) obtained in the previous section, the nuclear large subunit region (nrLSU), and the largest subunit of RNA polymerase II (RPB1), as well as the mitochondrial small subunit locus (mtSSU). The primers used for the amplification of these regions are provided in Supplementary Material Table 1.

PCR reactions were performed in a total volume of 15 μL, containing 2 μL of template DNA for amplification of the nuLSU, mtSSU (or 4 μL for the amplification of the RPB1 region), 0.5 μL of each primer (10 mM) and 6.5 μL of MyTaq Mix wich contains MyTaq DNA Polymerase (Bioline, UK)) and dNTPs; distilled water was added to reach the final volume.

The PCR conditions used for amplification for RPB1, nuLSU and mtSSU were the same as described in Perez-Ortega et al., (2016). PCR products were sequenced by Macrogen Inc. (Madrid, Spain) using the same primers as in the amplification reaction.

The four regions were aligned through MAFT tool v 1.4.0 (Katoh and Standley 2013) using the algorithm “auto” implemented in Geneious Prime v 2019.0.3 (https://www.geneious.com/prime/). To eliminate the ambiguously aligned regions of those alignments, Gblocks v 0.91b (http://molevol.cmima.csic.es/castresana/Gblocks_server.html) has been used (Castresana, 2000) allowing the least stringent parameters smaller final blocks, gap position within the blocks and less strict flanking positions. In order to select the best-fit partitioning schemes and nucleotide evolution models, we used PartitionFinder v 2.1.1 (Lanfear et al., 2017). Phylogenetic relationships among taxa were calculated using Bayesian inference in BEAST2 v 2.6 (Bouckaert et al., 2014) as implemented in CIPRES Science Gateway v 3 (https://www.phylo.org/) (Miller et al., 2010). We used an uncorrelated relaxed model with a log-normal prior for modelling clock and a Yule process to model tree prior (Drummond et al., 2006). The MCMC chain length was run during 5×10^9^ generations and results were logged every 5000 generations. Trace plots and effective sample sizes (ESS) were examined by TRACER v 1.7 (Rambaut et al., 2018). Finally, after discarding the first 25% sampled trees (burning), the results were summarized and annotated in a maximum clade credibility tree (MCC) through TreeAnnotator v 1.8.4 (https://beast.community/treeannotator) (Drummond et al., 2007). For the visualization of the resulting consensus tree and comparison of retrieved phylogenetic relationships with previous literature, we used Figtree v 1.4.4 (Rambaut, 2012). Accession numbers for the sequences generated and used during this study are available in Supplementary Material Table 2.

### Photobiont

A small fragment (1-2 mm^2^) of the thallus was taken from each specimen. Special care was taken to avoid thallus areas with clear signs of epiphytic fungi or algae as well as necrotic areas. Up to ten fragments per species and forest fragment were pooled in a single microcentrifuge tube. DNA extractions were performed using E.Z.N.A. Forensic DNA Kit (Omega Bio-Tek, Norcross, Georgia, USA) following the manufacturer’s instructions. The second part of the internal transcribed spacers (ITS2) of the photobiont was amplified using the primers FDGITS2-f y FDGITS2-r (Dal Grande et al., 2018) with Illumina adapters CS1 y CS2 attached (Available in: https://rtsf.natsci.msu.edu/-genomics/sequencing-services/sample-requirements-for-illumina-sequencing/). PCR reactions were performed using a mix of 25 μl which contained 0.625 U PrimerSTAR GXL DNA Polymerase (Takara BioInc, Japan), 5 μl of Buffer, 2 μl of dNTP Mixture (2.5 mM), 3 μl of DNA template, 0.5 μl of each primer at 10 mM and 13.5 μl of sterile distilled H_2_O. PCR conditions were set as follows: a 94°C denaturation step for 1 min followed by 30 cycles of 95°C for 15 s, 52°C for 15 s and 72 °C for 30 s ending with an elongation of 72°C for 1 min. PCR products were visualized in agarose gels and quantified by using a Qubit® Fluorometer (Life Technologies, Darmstadt, Germany) and normalized. Samples were indexed and pooled in a single Illumina MiSeq run (2 × 250bp paired-end sequencing, v2 Standard 500 cycle) at the Research Technology Support Facility Genomics Core at Michigan State University (USA).

### Bioinformatic analysis

MiSeq reads were merged, demultiplexed and filtered using the package DADA2 v 1.8.0 (https://benjjneb.github.io/dada2/index.html) for R studio (v 3.4.3 https://cran.r-project.org/bin/windows/base/old/3.4.3/). The first-left 22 bp corresponding to primer sequences were trimmed using the *filterAndTrim* function. Based on quality plots forward and reverse reads were trimmed to 200 and 170 bp respectively. Sequence filtering parameters was set to maxN = 0, maxEE=c (1,4), trunQ =2. The remaining parameters were operated as default. Finally, amplicon sequence variants (ASVs) were generated and a data matrix was built in which the rows represented the species of mycobiont, columns the ASVs of the photobiont and each cell contained the number of reads of each ASV inside the lichen, being the last a proxy for the interaction strength. The table containing the data was rarefied using the function “*rrarefy”* from the package “*vegan*”.

We could not rule out the presence of contaminations in spite of the special care taken during sample tissue collection, as thallus surface may dwell single algae cells not visible under the stereoscope. Further, some algal species may occur inside the thallus but not taking part in the symbiosis (Moya et al., 2017). In order to overcome this problem, we performed a double filtering of the obtained results. First of all, we collected information available in the literature about the photobionts associated with the species and/or genera found in our study (Blaha et al., 2006; Muggia et al., 2014; Tschermak-Woess, 1988; Wornik and Grube, 2010). Secondly, we established a limit of reads (100) below which we considered that the signal corresponded to contaminations of transients or algae not primarily associated with the mycobiont. Thus, sequences of taxa which based on the available literature do not represent the actual photobiont of the species and/or those with less than 100 reads were eliminated from subsequent analyses. All sequences obtained in this study will be available in the SRA (NCBI).

### Species specialization

An interaction matrix between mycobionts and photobionts was built to account for interactions in each forest patch. Mycobiont specialization towards their photobionts was explored by means of two metrics, normalized degree and the parameter *d’*. Normalized degree is the number of links (partners) per species normalized by the total number of possible partners. The parameter *d’* measures the specialization of each species based on its discrimination from a random selection of partners, ranging from 0 to 1 (a species interacts with only one other species, and that species only interacts with the first one) (Blüthgen et al., 2006). Both metrics were calculated for every species at all studied forest fragments using the function *specieslevel* from the *bipartite* v. 2.16 R package (Dormann et al., 2008).

### Statistical analyses

We first explored changes in ND and *d’* at the species level using Pearson correlation tests between each of the two specialization parameters and the fragment size gradient in those species present in at least five fragments. Analyses were carried out using the function *cor*.*test* of the *stats* v. 3.6.2 R package. Then, we explored the independent and cross effect of fragment size and two life traits, growth form and reproduction mode, on species specialization. All species recorded during the inventories fitted well within the main three growth forms recognized in lichens (Nash, 1996), i.e. crustose, whose thalli are strongly attached to the substrate and cannot be separated from it; foliose, whose thalli have a more or less flat appearance and are attached to the substratum by specialized structures in the lower cortex; and fruticose, whose thalli have a three-dimensional architecture, similar to tiny bushes and are attached to the substratum from a single point. Secondly, we categorized all species based on their reproduction modes. Although several reproductive strategies may be found in lichens (Tripp and Lendemer, 2018), species may be easily divided between those producing meiotic propagules (ascospores) which once settled in a new substrate they must find a compatible photobiont to reestablish the symbiosis, and those producing asexual propagules in which both partners, fungus and alga, are dispersed together, and the new thalli correspond to clones of the source thallus. We first explored the effect of area on the number of algae ASV and mycobiont species using the function *lm* of the package *stats*.

We investigated the effect of both functional traits, growth form and reproduction type in the mycobiont specialization towards their photosynthetic partners independently and in the context of a gradient of habitat availability. We used generalized linear mixed models in a Bayesian framework (Hadfield and Nakagawa, 2010) implemented in the package MCMCglmm v. 2.29 (Hadfield, 2010) in order to control the phylogenetic effects on these relationships. The uncertainty in the phylogenetic reconstructions was accounted for by running three MCMCglmm’s, each one using a phylogenetic tree randomly chosen from the distribution of tree topologies obtained in BEAST after discarding 25% of samples as burnin, and integrated over the posterior samples by drawing 1000 random samples across models and using *HPDinterval* function from the *coda* R package Plummer *et al*., 2013). Models priors follow de Villemereuil and Nakagawa (2014) using an inverse-Gamma distribution with shape and scale parameters equal to 0.01 as priors for the random effects and for the residual variance.

All graphs for the analyses performed were created with the R packages *ggplot2* (Wickham 2016), *gridExtra* (https://CRAN.R-project.org/package=gridExtraand) and *wesanderson* (https://CRAN.R-project.org/package=wesanderson).

### Interaction turnover

In order to shed light on the origin of changes in mycobiont specialization along the fragment size gradient, we analyzed the interaction beta diversity using the framework proposed by Poisot et al. (2012). This approach decomposes the total beta diversity of interactions (β_WN_) between two sites into the interaction dissimilarity due to species turnover species (β_ST_) and interaction dissimilarity when species occur in both sites but interaction not (β_OS_) (Poisot et al. 2012).

## RESULTS

### Mycobionts

We detected a total of 44 species of lichen-forming fungi in the study area (Sup. Mat. Table 2). 17 species were crustose, 19 foliose and 8 fruticose. Twenty species reproduced mainly sexually whereas 24 showed vegetative reproduction ((Sup. Mat. Table 2). Number of mycobiont species per forest fragment ranged from 9 at the 0.002 ha fragment to 28 at the 12.21 ha, with an average of 22 species per fragment (Table 1), with a distribution along the area that was adjusted to a quadratic model (R^2^= 0.82; p<0.05; Figure 2). The number of exclusive taxa was low in all fragments, occurring the highest, 2, at the largest forest fragment (Table 1).

**Figure 2.**
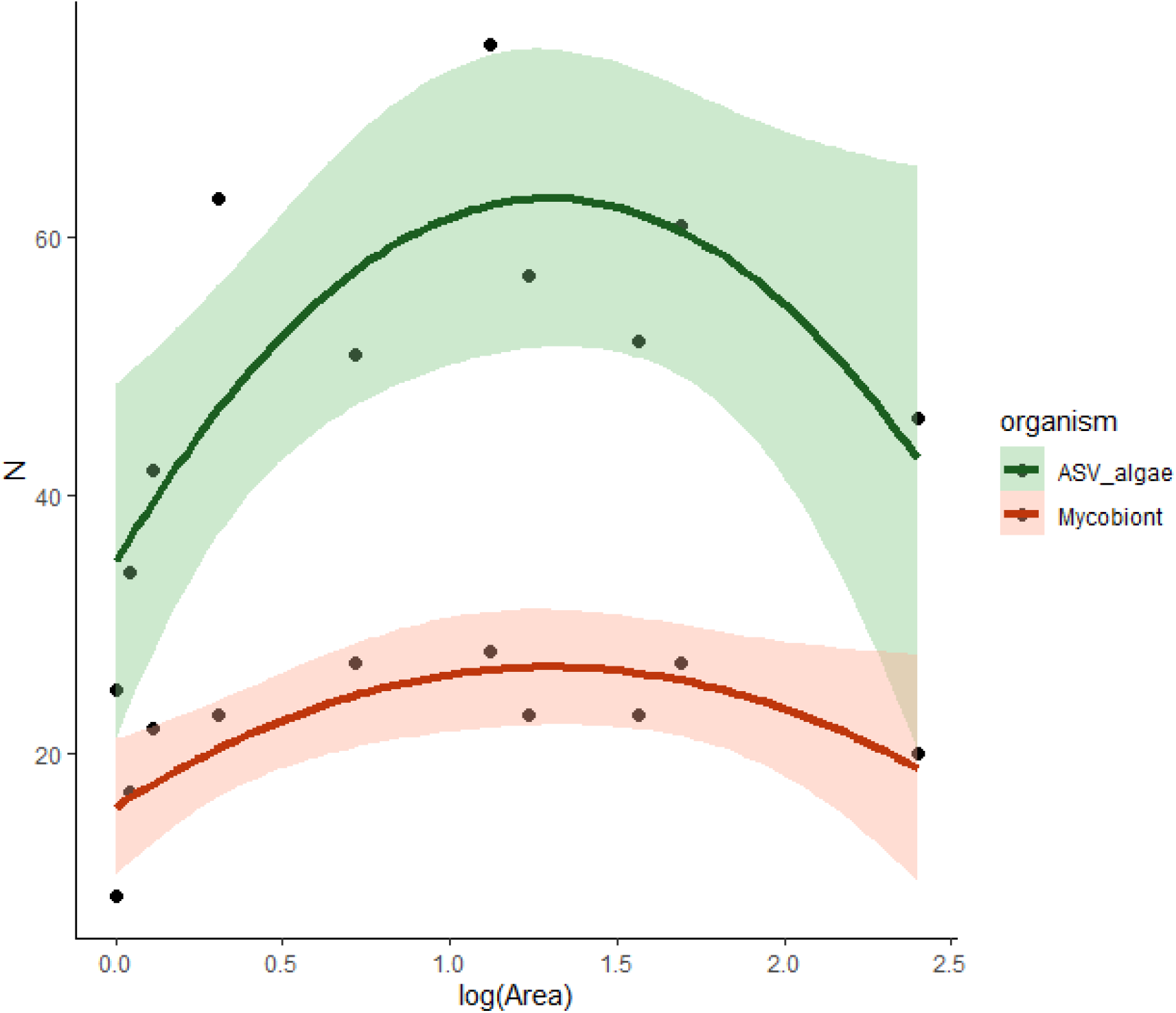
Distribution of number (N) of mycobiont species and ASVs along the patch size gradient (log(hectares)). Quadratic adjusted models (bold line) and confidence intervals (paler colored area) are represented for the number of mycobiont species in red (R^2^= 0.82; p<0.05) and green for ASVs (R^2^= 0.52; p <0.05).

The richness of crustose and foliose species did not vary along the area gradient while that of fruticose species showed a quadratic relation (R^2^= 0.571; F= 6.998, 2; d.f.=7; p-value<0.05), being maximum in intermediate plots. Regarding reproduction mode, the richness of sexual species showed a quadratic relationship with the log of area (R^2^= 0.617; F= 8.237, 2; d.f.=7; p-value<0.05), while the relationship for asexual species followed a positive linear trend (R^2^= 0.346; F= 5.756, 1; d.f.=8; p-value<0.05).

### Photobionts

A total of 2272 thalli were collected, corresponding to the 229 unique pools used for Illumina MiSeq analysis. The average number of thalli per species and fragment was 9.92. A total of 8,478,480 paired-end reads were obtained from the photobiont ITS2 amplicons. After quality filtering, a total of 4,956,878 reads remained. The processing of these reads resulted in 985 ASVs. After this processing we applied a manual filter removing the ASVs with less than 100 reads, obtaining a final number of 175 ASVs. Furthermore, the total number of reads remaining after this manual filter was 4,903,976 with an average of 21,415 reads per sample. BLAST searches of these remaining ASVs resulted in 158 corresponding to the genus *Trebouxia*, 13 to *Dictyochloropsis* and 4 to *Asterochloris. Asterochloris* ASVs were restricted to *Cladonia*, while the genus *Dictyochloropsis* was mostly found in the species *Phlyctis argena*.

The number of ASVs per forest fragment ranged from 25 at 0.002 ha to 75 at 12.21 ha (Table 1). ASV diversity along the patch size gradient followed a quadratic relationship (R^2^= 0.52; p <0.05; Figure 2). The highest number of exclusive photobiont ASVs (20) occurred at the 12.21 ha forest fragment (Table 1).

### Specialization

The mean number of ASVs per mycobiont species, species degree, ranged from 2.9 at the 250 ha fragment to 11.87 at the 16.13 ha fragment (Table 1). The normalized degree ranged from 0.01 in *Evernia prunastri* at the 12.21 ha forest fragment to 0.54 in *Blastenia xerothermica* at the 0.3 ha forest fragment (Supplementary Material Table 3). The specialization parameter *d’* ranged from 1 in *Cladonia fimbriata* at the 563.9 ha forest fragment to 0.16 in *Lecanora subcarpinea* at the 1.02 ha forest fragment (Supplementary Material Table 3).

The correlation analyses between ND and area in species with occurrences in ≥ 5 forest patches showed negative Pearson’s correlation coefficients for all species (Supplementary Material Table 4)., although they were only statistically significant in six of them, i.e. *Candelariella xanthostigma, Melanelixia subaurifera, Parmelina tiliacea, Physconia perisidiosa* and *Ramalina fastigiata*, all but the last one are vegetatively reproducing species. Regarding correlations between d’ and area, they were only significant in *Physconia enteroxantha* and *Physconia perisidiosa*, for which d’ increased along the forest fragment size gradient (Supplementary Material Table 4).

Phylogenetic generalized linear models showed differences in average normalized degree between crustose and fruticose species (Figure 3A), but not between reproduction modes (Figure 3B, Supplementary Material Table 4). Regarding the specialization parameter *d’* we found significant differences between crustose species and foliose and fruticose species (Figure 3C) as well as between reproduction modes, with vegetative species showing higher average d’ (Figure 3D, Supplementary Material Table 4).

**Figure 3.**
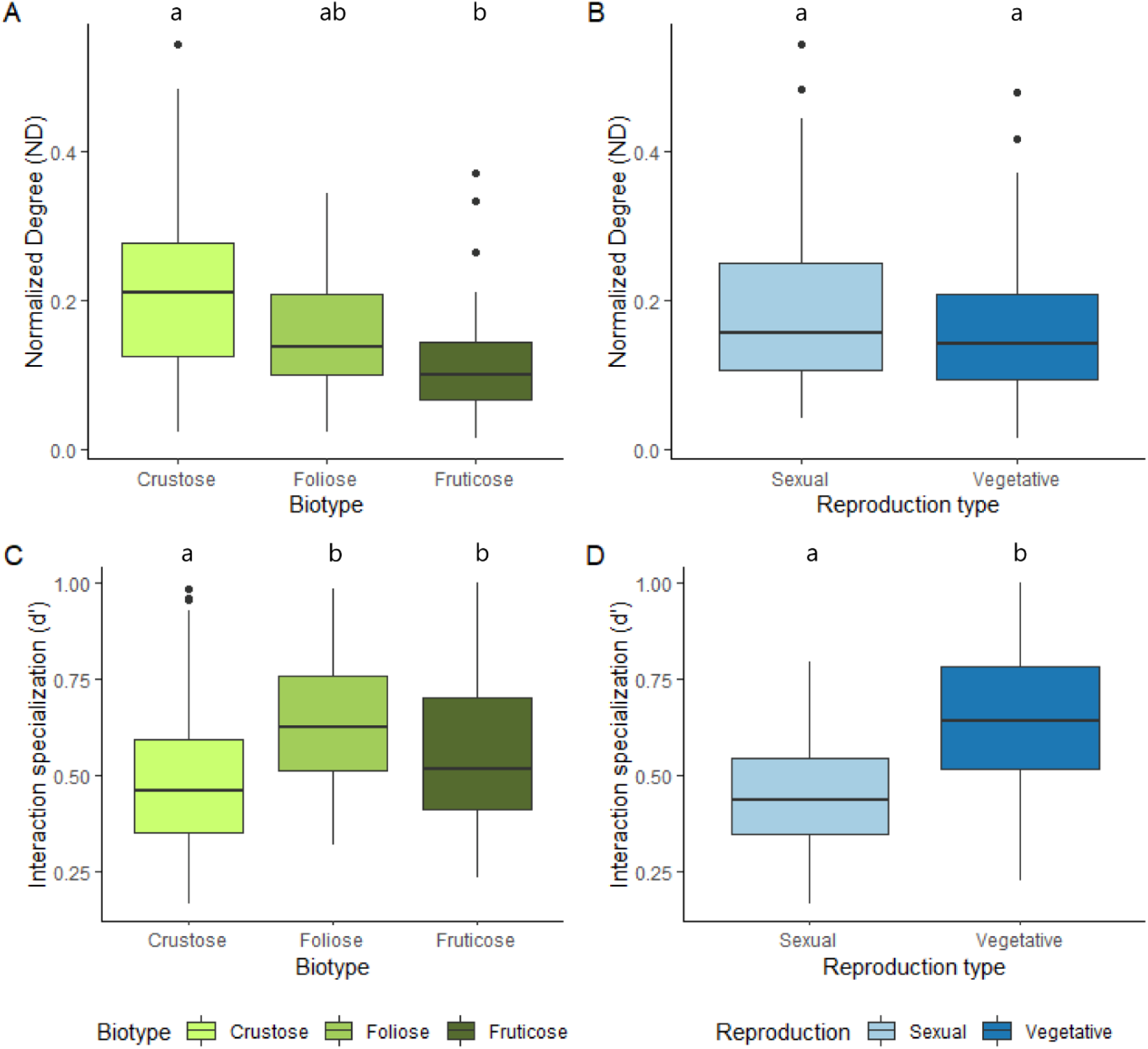
Boxplots show differences in specialization among species with different functional traits. A) Differences among biotypes in the specialization parameter *d’*; B) Differences between reproduction modes in the specialization parameter *d’*; C) Differences among biotypes in normalized degree (ND); D) Differences between reproduction modes in normalized degree (ND). The letters above boxplot depict significant differences among functional groups. Significance is based on phylogenetic controlled MCMCglm models (see. Mat. Supp. Table 3).

Phylogenetic generalized linear models also showed that area had a significant and negative effect on normalized degree (Figures 4A, B, Table 2). There were significant differences between crustose taxa and foliose and fruticose species and these differences were independent of the area (Table 2). No significant differences were found between reproduction modes regarding normalized degree along the area gradient (Table 2). Concerning the specialization parameter d’, the area had a significant effect, the larger the patch size the larger d’ when considering species biotypes (Figures 4C, D, Table 2). In addition, fruticose species had larger d’ than crustose species. Interestingly, there is a significant interaction between area and fruticose species, in which the increase of the parameter d’ along the area gradient is lower than in crustose species (Figure 4C, Table 2). When considering the effect of area and reproductive modes on d’ we found differences between sexual and asexual species, with significant effects of interactions (Figure 4D, Table 2). Vegetatively reproduced species increased their d’ along the area gradient, but the pattern was the contrary in sexual species.

**Figure 4.**
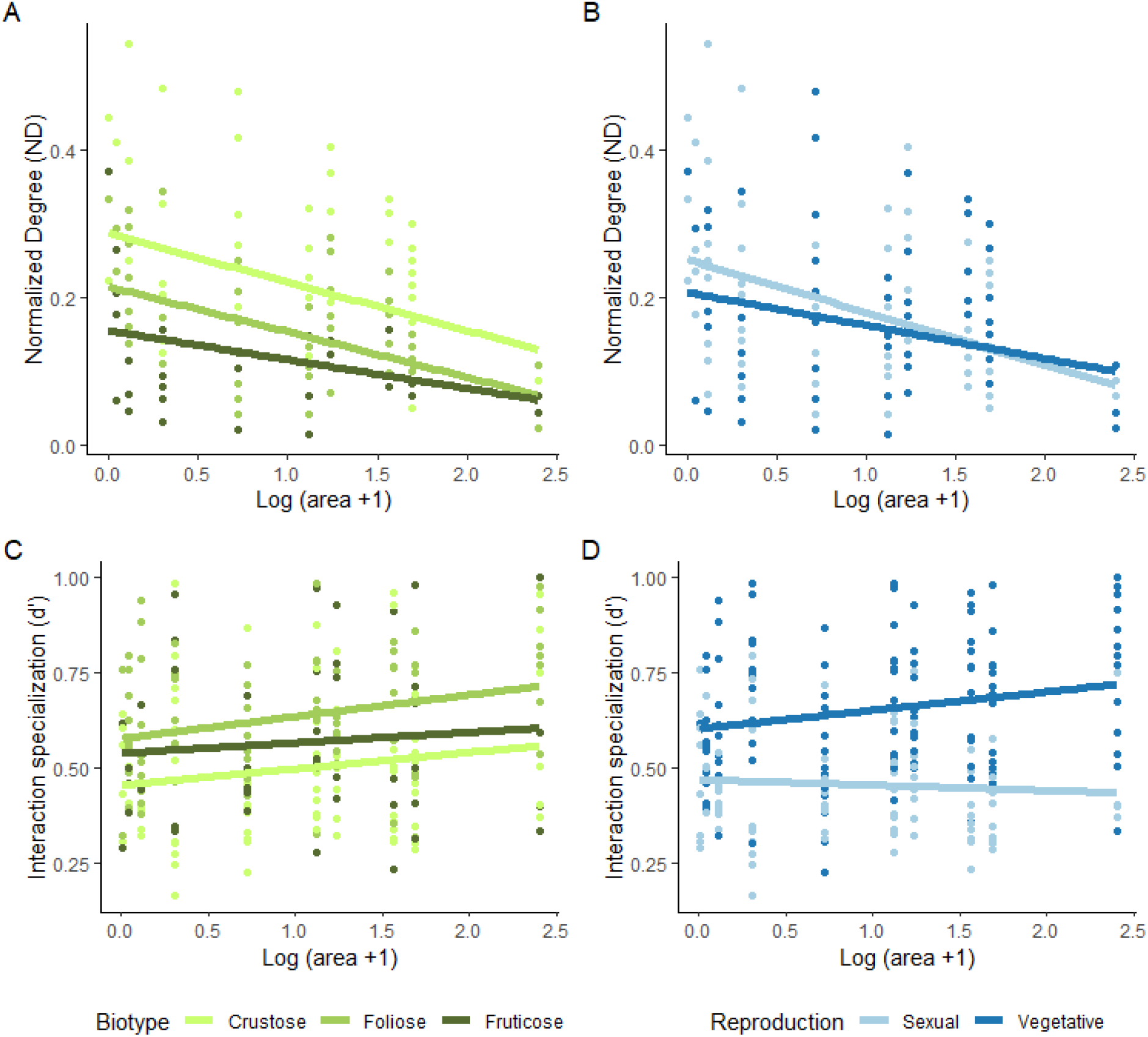
Representation of the interaction effect between area and reproduction type on ND (A) and d’(B), as well as the interaction effect between area and biotype on ND (C) and d’(D).

**Table 2.**
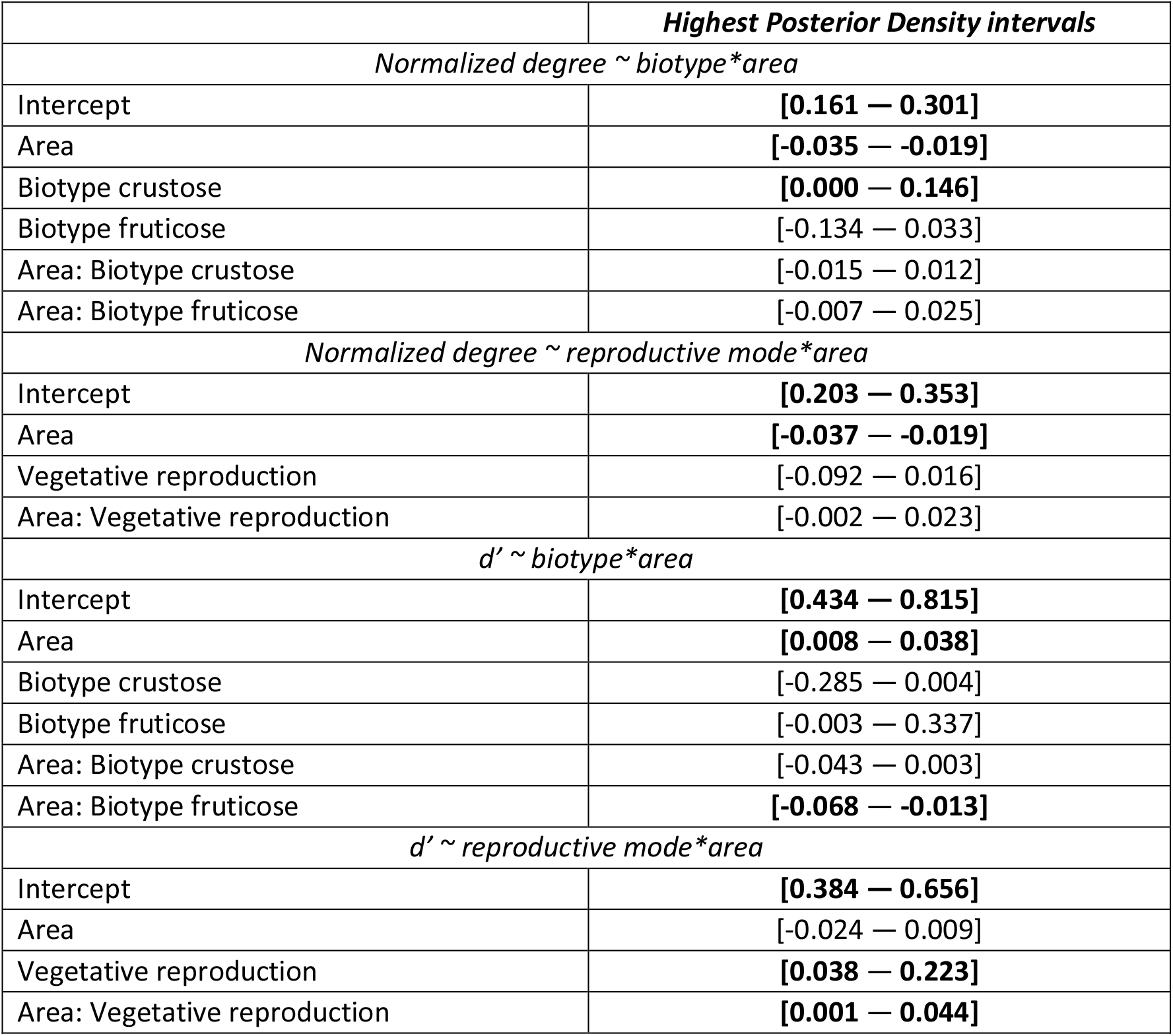
Results of Phylogenetic Generalized Linear Models explaining Normalized degree and specialization (d’) as a function of biotype, reproductive mode and area. Variables with significant effect, those in which the highest posterior density interval does not include zero, are shown in bold. Area was implemented in all analyses as log(area+1).

Interaction turnover analyses revealed that most of the differences between forest patches were due to the establishment of new interactions even though partners were present rather than dissimilarity due to partner turnover (β_ST_) (Figure. 5; t = 5.2741, df = 74.8, p-value<0.001).

**Figure 5.**
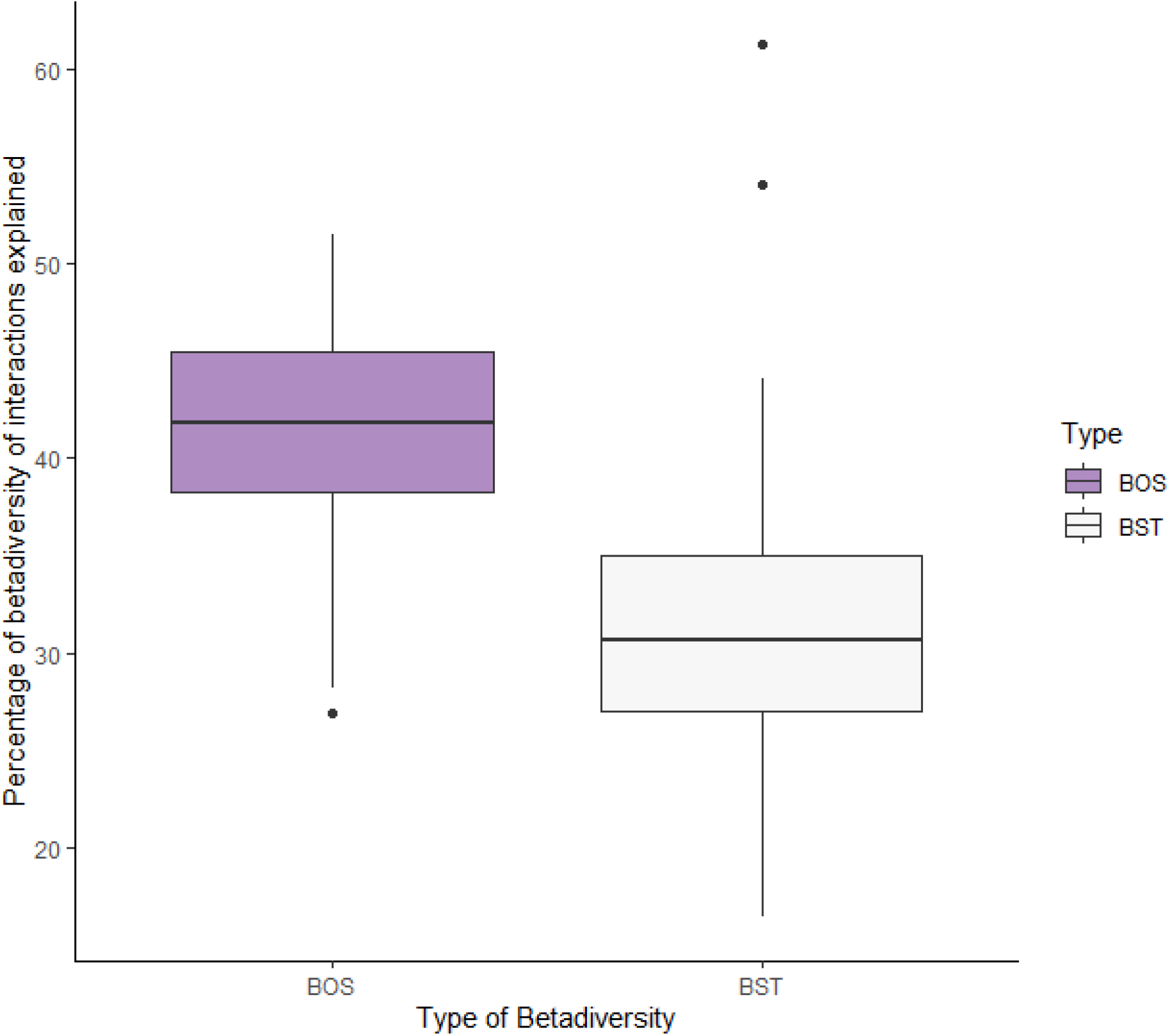
Percentage of beta diversity of interactions explained by the dissimilarity of interactions established between species common to both communities (BOS) and due to species turnover (BST) (t-test: t = 5.2741, df = 74.8, p-value<0.001).

## Discussion

Our study represents the first attempt to disentangle the effect of habitat loss on interaction specialization in symbiotic organisms. The obtained results shed light on how habitat loss interplays with life traits to affect fungal-algal interactions in epiphytic lichen symbioses.

### Diversity

Mycobionts and photobiont richness showed a humped-shaped distribution along the patch size gradient. This pattern is more pronounced for algae suggesting a higher sensitivity of algae to abiotic conditions. Previous studies dealing with lichen diversity along gradients of habitat loss in different forest ecosystems have reported a gradual loss of diversity as the available habitat diminished (Svoboda et al., 2010), with changes in abiotic conditions associated to an increasing fragment edge effect proposed as the main reason for such reduction (Asplund et al., 2014; Belinchón et al., 2007; Boudreault et al., 2008; Esseen and Renhorn, 1998). Hump-shaped diversity patterns have been reported for lichen-forming fungi along stand age (Asplund et al., 2014; Miller et al., 2020) and elevational gradients (Nanda et al., 2021) so-far this is the first report of a hump-shaped pattern of diversity along a fragmentation gradient. The intermediate disturbance hypothesis (IDH) (Connell, 1978) predicts highest levels of biodiversity at intermediate levels of frequency or intensity of disturbance (but see Mackey and Currie (2001) and Fox (2013) for criticism). Although it has been proposed that the main driver for this peak of diversity is the decrease of species densities, which entails reduced species competition (Fox 2013), it is not clear that this assumption may be applied to sessile organisms such as lichen-forming fungi with high habitat specialization (Fritz et al., 2009). It could be that fragments of intermediate size have intermediate abiotic conditions that allow the presence of habitat specialist populations as well as species associated with the abiotic conditions existing in smaller fragments. It is noteworthy that intermediate patch sizes dwell the largest number of exclusive ASVs, a pattern not shared by mycobionts. Therefore, the hump-shaped pattern found in mycobionts and photobionts may not respond to the same factors, or they may affect both organisms differently. Intermediate abiotic conditions could be translated into a higher range of available microhabitats, and higher spatial heterogeneity, promoting a larger amount of habitat specialists and exclusive ASVs in photobionts (Rosenzweig, 1995). If mycobionts had broader ecological requirements than photobionts, this would explain the difference in the observed pattern between the two organisms.

Our results showed that a gradient of habitat availability influences the distribution of functional traits in the studied lichen communities. Thus, except for crustose and foliose biotypes, the number of fruticose, vegetative and sexually reproduced species changed along the gradient. Changes in the distribution of functional traits in epiphytic lichen communities are directly related to abiotic conditions such as solar radiation, water availability or air pollution (Giordani et al., 2014; Koch et al., 2019; Matos et al. 2015; Paoli et al., 2017), and they change along environmental gradients such as elevation (Bässler et al., 2016) or land use (Pinho et al., 2012).

### Functional traits and specialization

Levels of specialization are affected by life traits in all known organisms (e.g. Armbruster, 2017; Dehling et al., 2016). Our results showed differences in specialization among biotypes and between reproductive modes, both in terms of ND and d’. Crustose species interacted on average with a higher number of ASVs than fruticose and foliose species, although only the first difference was significant. Regarding the parameter d’, crustose species showed significantly lower specialization than foliose and fruticose species. So far, no study had compared differences in specialization among biotypes although most studies on photobionts associated with crustose species have shown that they interact with a large number of species, in a wide phylogenetic spectrum (Blaha et al., 2006; Guzow-Krzeminska, 2006; Muggia et al., 2014) although examples of high specialization also occur (Hauck et al., 2007; Singh et al., 2015).

The interaction with a wider range of photobionts facilitates the settlement in a wider range of substrates and ecological conditions (Muggia et al., 2014) since changes in photobiont allow species to increase their ecological niche (Peksa and Škaloud, 2011; Fernández-Mendoza and Printzen, 2013; Rolshausen et al., 2018). However, why most crustose lichens studied so far show a higher number of associated photobionts still remains unclear. The constant contact with the substrate during lichen growth mediated by prothallus ensures the constant contact with new suitable free-living photobionts (Asmtrong and Bradwell, 2010; Sanders, 2005) that may be incorporated into the lichen thallus in the course of development (Sanders and Lücking, 2002). New photobionts might coexist with previous algal strains or replace them (Ott 1987). On the contrary, chances to incorporate new algal strains in foliose and fruticose species are much more reduced due to the scarce contact (single fixation points in fruticose species, rhizines in foliose not adnate species) with substrate and global isolation due to cortical tissues. Molins et al., (2021) has recently shown that early developmental stages of *R. farinacea* show higher algal diversity than older stages. This fact might indicate that, likewise it occurs in corals (Abrego et al., 2009); it is in early stages of foliose or fruticose species when there is more intimate contact with the substrate, therefore the mycobiont being able to establish contact with a wider array of algal strains, although only one or a few of these will remain in the later stages.

The reproduction mode also showed a clear effect on mycobiont specialization. Sexual and vegetatively reproduced species had on average a similar number of partners but the latter showed considerably higher values of d’. Contrary to our findings, previous studies comparing closely related species pairs had shown that species of lichen-forming fungi with clonal reproduction via joint dispersal propagules tend to have lower associated algal diversity and are more selective than sexual species (Cao et al., 2015; Otálora et al., 2013). The lower diversity associated with species that reproduce by vegetative propagules has been attributed to the joint dispersal of both bionts and the lack of need to search for new partners if the fitness of the combination is optimal or suboptimal in the new environment. However, this interaction may be broken and the mycobiont may interact with new photobionts in the new substrate, using the original photobiont only for the settlement stage, thus avoiding the restriction to colonize new environments. So far, no study had analyzed a high number of species to have a broad overview. Thus, we showed that, on average, sexual and asexual species did not differ in terms of the number of associated photosynthetic partners. Wornik and Grube (2010) showed that the ease to switch photobiont in the vegetatively reproduced *Physconia grisea*, a Mediterranean species dispersed via soredia and also found in our study area, was the only suitable explanation for comparable photobiont diversity between a sexual and an asexual species. However, regarding d’ differences between both reproductive modes were significant and outstanding, showing asexual species much higher specialization. Cao et al. (2015) suggested that the decrease of selectivity in sexual species may be due to the need of finding a new photobiont before the ascospores perish. In this case, the similar number of partners in both groups does not point in that direction. The greater specialization found in vegetatively dispersing species may reflect the higher fitness of a particular myco-photobiont combination at the local scale which is favored by the clonal reproduction.

### Specialization along the gradient of area size

Our results showed that average specialization at the species level changed along the gradient of fragment size in the same way as occurs with other parameters that describe the structure of interaction networks (Tylianakis and Morris, 2017). The variation of species specialization in such gradients has only been scarcely studied (Aizen et al., 2012). Specialization is often considered a species-level trait, conserved across the phylogeny (Gómez et al., 2010) and with relatively little variation, although it is known to vary at geographical and phylogenetic scales (Poisot et al., 2015; Trøjelsgaard et al., 2015). Carstensen et al. (2018) found that local and regional specialization were strongly correlated in 14 plant-pollinator networks from two different continents, considering species specialization consistent level across scales. On the contrary, we found that average specialization may change due to habitat abiotic conditions, here derived from habitat fragmentation. We observed a strong decrease in the average number of photobionts associated with each fungal species as the fragment size decreased and an overall higher specialization (*d’*) along the area gradient. The higher values of the normalized degree found in smaller patch sizes regarding larger ones could be an artifact derived from the lower number of photobionts available in small fragments (Fig. 2). However, the specialization parameter *d’* did clearly point towards higher mycobiont specialization in larger fragments. Although the number of available partners increased along most of the gradient, fungal species become more selective. In addition, the analysis of the interaction beta diversity showed that most of the turnover among forest patches was not due to changes in biont composition, that is, the observed pattern of increasing specialization is not due to the absence of certain bionts, but to an increase in the level of selectivity among the those available. Assuming a local adaptation of the photobiont, the smaller fragments, with more homogeneous abiotic conditions due to the greater edge effect (greater insolation, wind, nitrogen inputs, etc.) may host only photobiont species that we could call weeds, adapted to the greater intensity of the disturbance. New propagules arriving in the patch will have at their disposal only a pool of equally suboptimal photobionts, perhaps interchangeable in terms of holobiont fitness, so the degree of selectivity is low. On the contrary, in larger fragments, where a larger pool of photobionts is available but also abiotic conditions are more heterogenous propagules of different species will be able to establish more specialized interactions, thus increasing holobiont fitness.

### Interplay between specialization, traits and fragmentation

Changes in specialization along the area gradient were not even among morphological groups and between the reproduction modes, showing that changes in specialization along the gradient are also influenced by their life traits. For example, although the number of interacting photobionts decreases along the area gradient in all biotypes, there was a significant difference between how this number decreases in foliose and fruticose species and how it does in crustose species. In addition, we found a significant interplay between the fruticose biotype and area, regarding the specialization parameter d’, showing that changes in habitat conditions and availability do not affect similarly all biotypes in terms of discriminating among their photosynthetic partners. It is known that different growth forms represent different strategies of exploitation of water sources in lichens (Gauslaa, 2014). Thus, it is likely that the changing abiotic conditions along the patch size gradient do not affect similarly photobionts occurring in different growth forms since thallus biotypes may have different buffering potential of environmental conditions.

Regarding reproductive modes, it is certainly striking that although the ND of both functional groups similarly decreased along the area gradient, they showed contrasting trends regarding the specialization parameter d’ with sexual species decreasing in their specialization as the patch area increases. The mode of reproduction has relevant implications with respect to constraints on dispersal and colonization of new habitats (Ellis et al., 2021; Walser, 2004), thallus physiology (Tretiach et al., 2005), and response to disturbance gradients such as air pollution (Koch et al., 2019). The contrasting trends along the area gradient between reproductive modes in the average d’ may have their origin in differences in species composition and the distribution of reproductive modes in the communities and also, in the different modulation of specialization along the gradient at the individual level. Models indicated a significant change of reproduction modes along the fragment size area. In addition, linear models for individual changes in specialization along the gradient revealed five species in which species significantly modulated their specialization in terms of ND (four of them asexually reproduced via soredia) and two cases in which d’ significantly increased along the gradient, both asexually reproduced via soredia. Thus, the explanation of this contrasting pattern may be derived both from changes in species composition and modulation of specialization in certain species. Modulation of specialization at the species level is certainly a phenomenon that should be further explored in the future.

### Implications for conservation

Our results have clear implications in conservation biology. Interactions among species are certainly influenced by local abiotic conditions, so a certain biont combination would not have the same fitness in two patches with different sizes. Small changes in forest conditions can have drastic consequences and cause existing biont combinations to lose their ability to compete. In addition, if some species can modulate their specialization to a certain degree, then those species would display higher resilience to environmental changes such as those derived from habitat fragmentation than other species. Their ability to adapt to changes in local conditions through changes in specialization towards their photobionts may buffer the effect of more stressful abiotic conditions due to the increase of edge effect in small fragments, at least until phylogenetic constraints prevent interactions in some species. This buffering effect produced by the modulation of specialization may postpone the extinction debt (Berglund and Jonsson, 2005), or, depending on the intensity and frequency of perturbation, even avoid extinction, with remarkable implications for biodiversity conservation (Kuussaari et al., 2009). Further studies will show whether the analyses of interaction dynamics (partner fidelity, link conservatism and rewiring) in myco-photobiont symbioses in the context of interaction bipartite networks may be used as suitable tools to understand and predict long-term changes in epiphyte lichen communities.

## Author contributions

SPO, MV, MVdP and ABdG; SPO and ABdG designed the study; SPO and ABdG carried out the field sampling; ABdG and MVdP collected laboratory data; MV, SPO, ABdG, MVdP and MV analyzed the data; SPO and ABdG led the writing of the manuscript; All authors contributed critically to the drafts and gave final approval for publication.

## Data availability statement

All NGS sequences obtained in this study will be available in the SRA (NCBI). Sequences obtained through Sanger sequencing are available in Gen Bank (Accession numbers provided in Supplementary Material Table 2).

## Acknowledgments

This study was partly funded by the grant PID2019-111527GB-I00 from the Spanish Ministry of Science and Innovation. SPO was supported by the grant RYC-2014-16 784 from the Spanish Ministry of Economy, Industry and Competitiveness.

## REFERENCES

Abrego, D., Van Oppen, M.J.H., & Willis B. (2009). Onset of algal endosymbiont specificity varies among closely related species of Acropora corals during early ontogeny Molecular Ecology 18:3532–3543 doi:https://doi.org/10.1111/j.1365-294X.2009.04276.x

Aizen, M., Sabatino, M., & Tylianakis, J. (2012). Specialization and Rarity Predict Nonrandom Loss of Interactions from Mutualist Networks. Science, vol 335. doi:10.1126/science.1215320

Aragón, G., Martínez, I., Izquierdo, P., Belinchón, R., & Escudero, A. (2010). Effects of forest management on epiphytic lichen diversity in Mediterranean forests Applied Vegetation. Science, 13:183–194 doi:10.1111/j.1654-109X.2009.01060.x

Armbruster, W.S. (2017). The specialization continuum in pollination systems: diversity of concepts and implications for ecology, evolution and conservation. Functional Ecology, 31:88–100 doi:https://doi.org/10.1111/1365-2435.12783

Asmtrong, R., & Bradwell, T. (2010). Growth of crustose lichens: A review Geografiska Annaler: Series A, Physical Geography 92:3–17 doi:https://doi.org/10.1111/j.1468-0459.2010.00374.x

Asplund, J., Sandling, A., Kardol, P., & Wardle, D.A. (2014). The influence of tree-scale and ecosystem-scale factors on epiphytic lichen communities across a long-term retrogressive chronosequence. Journal of Vegetation Science, 25:1100–1111 doi:https://doi.org/10.1111/jvs.12149

Asta, J., Erhardt, W., Ferretti, M., Fornasier, F., Kirschbaum, U., Nimis, P. L., Purvis, O.W., Pirintsos, S., Scheidegger, C., Van Haluwyn, C., & Wirth, V. (2002).Mapping lichen diversity as an indicator of environmental quality In: Nimis P.L., Scheidegger C., Wolseley P.A. (eds) Monitoring with Lichens — Monitoring Lichens. NATO Science Series (Series IV: Earth and Environmental Sciences), vol 7. Springer, Dordrecht. https://doi.org/10.1007/978-94-010-0423-7_19

Bagchi, R., Brown, L.M., Elphick, C.S., Wagner, D.L., & Singer, M.S. (2018). Anthropogenic fragmentation of landscapes: mechanisms for eroding the specificity of plant– herbivore interactions. Oecologia, 187:521–533 doi:10.1007/s00442-018-4115-5

Bässler, C., Cadotte, M. W., Beudert, B., Heibl, C., Blaschke, M., Bradtka, J. H., Langbehn, T., Werth, S., Müller, J. (2016). Contrasting patterns of lichen functional diversity and species richness across an elevation gradient. Ecography, 39:689–698 doi: https://doi.org/10.1111/ecog.01789

Belinchón, R., Martínez, I., Escudero, A., Aragón, G., & Valladares, F. (2007). Edge effects on epiphytic communities in a Mediterranean Quercus pyrenaica forest. Journal of Vegetation Science, 18:81–90 doi:https://doi.org/10.1111/j.1654-1103.2007.tb02518.x

Berglund, H., & Jonsson, B.G. (2005). Verifying an Extinction Debt among Lichens and Fungi in Northern Swedish Boreal Forests. Conservation Biology, 19:338–348 doi:https://doi.org/10.1111/j.1523-1739.2005.00550.x

Blaha, J., Baloch, E., & Grube, M. (2006). High photobiont diversity associated with the euryoecious lichen-forming ascomycete Lecanora rupicola (Lecanoraceae, Ascomycota). Biological Journal of the Linnean Society, 88:283–293 doi:10.1111/j.1095-8312.2006.00640.x

Blüthgen, N., Menzel, F., & Blüthgen, N. (2006). Measuring specialization in species interaction networks. BMC Ecology, 6:9 doi:10.1186/1472-6785-6-9

Boudreault, C., Bergeron, Y., Drapeau, P., & Mascarúa López. L. (2008). Edge effects on epiphytic lichens in remnant stands of managed landscapes in the eastern boreal forest of Canada. Forest Ecology and Management, 255:1461–1471 doi:10.1016/j.foreco.2007.11.002

Bouckaert, R., Heled, J., Kühnert, D., Vaughan, T., Wu, C. H., Xie, D., Marc, A., Suchard, A.R., Rambaut, A., & Drummond, A. J. (2014). BEAST 2: a software platform for Bayesian evolutionary analysis. PLoS Comput Biol, 10(4), e1003537. https://doi.org/10.1371/journal.pcbi.1003537

Bovo, A.A.A., Ferraz, K.M.P.M.B., Magioli, M., Alexandrino, E.R., Hasui, É., Ribeiro, M.C., & Tobias, J.A. (2018). Habitat fragmentation narrows the distribution of avian functional traits associated with seed dispersal in tropical forest. Perspectives in Ecology and Conservation, 16:90–96 doi:https://doi.org/10.1016/j.pecon.2018.03.004

Brunialti, G., Frati, L., & Loppi, S. (2013). Fragmentation of Mediterranean oak forests affects the diversity of epiphytic lichens. Nova Hedwigia, 96:265–278. doi: 10.1127/0029-5035/2012/0075

Buschbom, J., & Mueller, G.M. (2005). Testing “species pair” hypotheses: evolutionary processes in the lichen-forming species complex Porpidia flavocoerulescens and Porpidia melinodes. Molecular Biology and Evolution, 23:574–586. https://doi.org/10.1093/molbev/msj063

Cao, S., Zhang, F., Liu, C., Hao, Z., Tian, Y., Zhu, L., & Zhou, Q. (2015). Distribution patterns of haplotypes for symbionts from Umbilicaria esculenta and U. muehlenbergii reflect the importance of reproductive strategy in shaping population genetic structure. BMC Microbiology, 15:212 doi:10.1186/s12866-015-0527-0

Carstensen, D.W., Trøjelsgaard, K., Ollerton, J., & Morellato, L.P.C. (2018). Local and regional specialization in plant–pollinator networks. Oikos, 127:531–537 doi:https://doi.org/10.1111/oik.04436

Castresana, J. (2000). Selection of conserved blocks from multiple alignments for their use in phylogenetic analysis. Molecular biology and evolution, 17(4), 540–552. https://doi.org/10.1093/oxfordjournals.molbev.a026334

Connell, J.H. (1978). Diversity in Tropical Rain Forests and Coral Reefs. Science, 199:1302–1310 doi:doi:10.1126/science.199.4335.1302

Dal Grande, F., Rolshausen, G., Divakar, P.K., Crespo, A., Otte, J., Schleuning, M., & Schmitt, I. (2018). Environment and host identity structure communities of green algal symbionts in lichens. New Phytologist, 217:277–289 doi:10.1111/nph.14770

Dal Grande, F., Widmer, I., Wagner, H.H., & Scheidegger, C. (2012). Vertical and horizontal photobiont transmission within populations of a lichen symbiosis. Molecular Ecology, 21:3159–3172 doi:10.1111/j.1365-294X.2012.05482.x

de Villemereuil, P., & Nakagawa, S. (2014). General Quantitative Genetic Methods for Comparative Biology. In: Garamszegi LZ (ed) Modern Phylogenetic Comparative Methods and Their Application in Evolutionary Biology: Concepts and Practice. Springer Berlin Heidelberg, Berlin, Heidelberg, pp 287–303. doi:10.1007/978-3-662-43550-2_11

Dehling, D.M., Jordano, P., Schaefer, H.M., Böhning-Gaese, K., & Schleuning, M. (2016). Morphology predicts species’ functional roles and their degree of specialization in plant-frugivore interactions. Proceedings of the Royal Society, B: Biological Sciences 283:20152444 doi:10.1098/rspb.2015.2444

Dennis, R.L.H., Dapporto, L., Fattorini, S., & Cook, L.M. (2011). The generalism–specialism debate: the role of generalists in the life and death of species. Biological Journal of the Linnean Society, 104:725–737 doi:10.1111/j.1095-8312.2011.01789.x

Devictor, V., Julliard, R., & Jiguet, F. (2008). Distribution of specialist and generalist species along spatial gradients of habitat disturbance and fragmentation. Oikos, 117:507–514 https://doi.org/10.1111/j.0030-1299.2008.16215.x

Dormann, C.F., Gruber, B., & Fründ, J. (2008). Introducing the bipartite package: analysing ecological networks interaction 1:0.2413793

Drummond, A. J., Ho, S. Y., Phillips, M. J., & Rambaut, A. (2006). Relaxed phylogenetics and dating with confidence. PLoS Biol, 4(5), https://doi.org/10.1371/journal.pbio.0040088

Drummond, A. J., & Rambaut, A. (2007). BEAST: Bayesian evolutionary analysis by sampling trees. BMC evolutionary biology, 7(1), 1–8. https://doi.org/10.1186/1471-2148-7-214

Ellis CJ et al., (2021). Functional Traits in Lichen Ecology: A Review of Challenge and Opportunity. Microorganisms, 9:766, https://doi.org/10.3390/microorganisms9040766

Esseen, P-A., & Renhorn, K-E (1998). Edge Effects on an Epiphytic Lichen in Fragmented Forests. Conservation Biology, 12:1307–1317 doi:https://doi.org/10.1111/j.1523-1739.1998.97346.x

Fahrig, L. (2003). Effects of Habitat Fragmentation on Biodiversity Annual Review of Ecology, Evolution, and Systematics, 34:487–515 doi:10.1146/annurev.ecolsys.34.011802.132419

Farneda, F.Z., Rocha, R., López-Baucells, A., Groenenberg, M., Silva, I., Palmeirim, J.M., Bobrowiec, P.E.D. & Meyer, C.F.J. (2015). Trait-related responses to habitat fragmentation in Amazonian bats. Journal of Applied Ecology, 52: 1381–1391. https://doi.org/10.1111/1365-2664.12490

Ferencova, Z., Rico, V.J, & Hawksworth, D.L. (2017). Extraction of DNA from lichen-forming and lichenicolous fungi: a low-cost fast protocol using Chelex. The Lichenologist, 49:521–525 doi:10.1017/s0024282917000329

Fernández-Mendoza, F., Printzen, C. (2013). Pleistocene expansion of the bipolar lichen Cetraria aculeata into the Southern hemisphere. Molecular Ecology, 22:1961–1983 doi:https://doi.org/10.1111/mec.12210

Fernández-Mendoza, F., Domaschke, S., García, M., Jordan, P., Martín, M.P., & Printzen, C. (2011). Population structure of mycobionts and photobionts of the widespread lichen Cetraria aculeata. Molecular Ecology, 20:1208–1232, https://doi.org/10.1111/j.1365-294X.2010.04993.x

Fox, J.W. (2013). The intermediate disturbance hypothesis should be abandoned Trends in Ecology & Evolution 28:86–92 doi:https://doi.org/10.1016/j.tree.2012.08.014

Fritz, Ö., Niklasson, M., & Churski, M. (2009). Tree age is a key factor for the conservation of epiphytic lichens and bryophytes in beech forests. Applied Vegetation Science, 12:93–106 doi:https://doi.org/10.1111/j.1654-109X.2009.01007.x

Gardes, M., & Bruns, T. D. (1993). ITS primers with enhanced specificity for basidiomycetes-application to the identification of mycorrhizae and rusts. Molecular Ecology, 2:113–118. doi: 10.1111/j.1365-294x.1993.tb00005.x.

Gauslaa, Y. (2014). Rain, dew, and humid air as drivers of morphology, function and spatial distribution in epiphytic lichens. The Lichenologist, 46:1–16 doi: https://doi.org/10.1017/S0024282913000753

Giordani, P., Brunialti, G., Bacaro, G., Nascimbene, J. (2012). Functional traits of epiphytic lichens as potential indicators of environmental conditions in forest ecosystems. Ecological Indicators, 18:413–420 https://doi.org/10.1016/j.ecolind.2011.12.006

Giordani, P., Incerti, G., Rizzi, G., Rellini, I., Nimis, P.L., & Modenesi, P. (2014). Functional traits of cryptogams in Mediterranean ecosystems are driven by water, light and substrate interactions. Journal of Vegetation Science, 25:778–792 doi:https://doi.org/10.1111/jvs.12119

Giordani, P., Calatayud, V., Stofer, S., Seidling, W., Granke, O., Fischer, R. (2014). Detecting the nitrogen critical loads on European forests by means of epiphytic lichens. A signal-to-noise evaluation. Forest Ecology and Management, 311:29–40 doi: 10.1016/j.foreco.2013.05.048.

Gómez, J.P., Bravo, G.A., Brumfield, R.T., Tello, J.G., & Cadena, C.D. (2010). A phylogenetic approach to disentangling the role of competition and habitat filtering in community assembly of Neotropical forest birds. Journal of Animal Ecology, 79:1181–1192 doi:https://doi.org/10.1111/j.1365-2656.2010.01725.x

Gonzalez, A., Rayfield, B., & Lindo, Z. (2011). The disentangled bank: How loss of habitat fragments and disassembles ecological networks. American Journal of Botany, 98:503–516 doi:https://doi.org/10.3732/ajb.1000424

Grube, M., Hawksworth, D.L. (2007). Trouble with lichen: the re-evaluation and re-interpretation of thallus form and fruit body types in the molecular era. Mycological Research, 111:1116–1132 doi:https://doi.org/10.1016/j.mycres.2007.04.008

Guzow-Krzeminska, B. (2006). Photobiont flexibility in the lichen Protoparmeliopsis muralis as revealed by ITS rDNA analyses. The Lichenologist, 38:469–476 doi:10.1017/S0024282906005068

Haddad, N. M., Brudvig, L. A., Clobert, J., Davies, K. F., Gonzalez, A., Holt, R. D., Lovejoy, T.E., Sexton, J.O., Austin, M.P., Collins, C.D., Cook, W.M., Damschen, E.I., Ewers, R.M., … & Townshend, J. R. (2015). Habitat fragmentation and its lasting impact on Earth’s ecosystems. Science advances, 1(2), e1500052. doi: 10.1126/sciadv.1500052

Hadfield, J. D. (2010). MCMC Methods for Multi-Response Generalized Linear Mixed Models: The MCMCglmm R Package. Journal of Statistical Software, 33(2), 1–22. https://doi.org/10.18637/jss.v033.i02

Hadfield, J. D., & Nakagawa, S. (2010). General quantitative genetic methods for comparative biology: phylogenies, taxonomies and multi-trait models for continuous and categorical characters. Journal of Evolutionary Biology, 23: 494–508. https://doi.org/10.1111/j.1420-9101.2009.01915.x

Hadley, A.S., Frey, S.J.K., Robinson, W.D., & Betts, M.G. (2018). Forest fragmentation and loss reduce richness, availability, and specialization in tropical hummingbird communities. Biotropica, 50:74–83 doi:https://doi.org/10.1111/btp.12487

Hauck, M., Helms, G., & Friedl, T. (2007). Photobiont selectivity in the epiphytic lichens Hypogymnia physodes and Lecanora conizaeoides. The Lichenologist, 39:195–204 doi:10.1017/S0024282907006639

Hawksworth, D.L., & Grube, M. (2020). Lichens redefined as complex ecosystems. New Phytologist, 227:1281–1283 doi:https://doi.org/10.1111/nph.16630

Henle, K., Davies, K.F., Kleyer, M., Margules, C., & Settele, J. (2004). Predictors of species sensitivity to fragmentation. Biodiversity & Conservation, 13:207–251 https://doi.org/10.1023/B:BIOC.0000004319.91643.9e

Honegger, R. (2001). The Symbiotic Phenotype of Lichen-Forming Ascomycetes. In: Hock B. (eds) Fungal Associations. The Mycota (A Comprehensive Treatise on Fungi as Experimental Systems for Basic and Applied Research), vol 9. Springer, Berlin, Heidelberg. https://doi.org/10.1007/978-3-662-07334-6_10***

Hurtado, P., Prieto, M., Martínez-Vilalta, J., Giordani, P., Aragón, G., López-Angulo, J., Košuthová, A., Merinero, S., Díaz-Peña, E. M., Rosas, T., Benesperi, R., Bianchi, E., Grube, M., Mayrhofer, H., Nascimbene, J., Wedin, M., Westberg, M., & Martínez, I. (2020). Disentangling functional trait variation and covariation in epiphytic lichens along a continent-wide latitudinal gradient. Proceedings of the Royal Society B,2872019286220192862 http://doi.org/10.1098/rspb.2019.2862

Isbell, F., Calcagno, V., Hector, A., Connolly, J., Harpole, W. S., Reich, P. B., Scherer-Lorenzen, M., Schmid, B., Tilman, D., van Ruiijven, J.,Weigelt, A., Wilsey, B.J., Zavaleta, E.S., & Loreau, M. (2011). High plant diversity is needed to maintain ecosystem services. Nature, 477(7363), 199–202. https://doi.org/10.1038/nature10282

Jauker, F., Jauker, B., Grass, I., Steffan-Dewenter, I., & Wolters, V. (2019). Partitioning wild bee and hoverfly contributions to plant–pollinator network structure in fragmented habitats. Ecology, 100:e02569 doi:https://doi.org/10.1002/ecy.2569

Katoh, K., & Standley, D.M. (2013). MAFFT Multiple Sequence Alignment Software Version 7: Improvements in Performance and Usability. Molecular Biology and Evolution, 30:772–780 doi:10.1093/molbev/mst010

Koch, N.M., Matos, P., Branquinho, C., Pinho, P., Lucheta, F., Martins, S.M.dA., Vargas, & V.M.F. (2019). Selecting lichen functional traits as ecological indicators of the effects of urban environment. Science of The Total Environment, 654:705–713 doi:https://doi.org/10.1016/j.scitotenv.2018.11.107

Kuussaari, M., Bommarco, R., Heikkinen, R. K., Helm, A., Krauss, J., Lindborg, R., Öckinger, E., Pärtel, M., Pino, J., Rodà, F., Stefanescu, C., Teder, T., Zobel, M., & Steffan-Dewenter, I (2009). Extinction debt: a challenge for biodiversity conservation. Trends in ecology & evolution, 24(10), 564–571. https://doi.org/10.1016/j.tree.2009.04.011

Lanfear, R., Frandsen, P. B., Wright, A. M., Senfeld, T., & Calcott, B. (2017). PartitionFinder 2: new methods for selecting partitioned models of evolution for molecular and morphological phylogenetic analyses. Molecular biology and evolution, 34(3), 772–773. https://doi.org/10.1093/molbev/msw260

Leavitt, S.D., Kraichak, E., Nelsen, M.P., Altermann, S., Divakar, P.K., Alors, D., Esslinger, T.L., Crespo, A., & Lumbsch, T. (2015). Fungal specificity and selectivity for algae play a major role in determining lichen partnerships across diverse ecogeographic regions in the lichen-forming family Parmeliaceae (Ascomycota). Molecular Ecology, 24: 3779–3797. https://doi.org/10.1111/mec.13271

Mackey, R.L., & Currie, D.J. (2001). The diversity–disturbance relationship: is it generally strong and peaked?. Ecology, 82:3479–3492 doi:https://doi.org/10.1890/0012-9658(2001)082[3479:TDDRII]2.0.CO;2

Magain, N., Miadlikowska, J., Goffinet, B., Sérusiaux, E., & Lutzoni, F. (2016). Macroevolution of specificity in cyanolichens of the genus Peltigera section Polydactylon (Lecanoromycetes, Ascomycota). Systematic biology, 66:74–99 https://doi.org/10.1093/sysbio/syw065

Maglianesi, M.A., Blüthgen, N., Böhning-Gaese, K., & Schleuning, M. (2014). Morphological traits determine specialization and resource use in plant–hummingbird networks in the neotropics. Ecology, 95:3325–3334 doi:10.1890/13-2261.1

Magrach, A., Laurance, W.F., Larrinaga, A.R., & Santamaria, L. (2014). Meta-analysis of the effects of forest fragmentation on interspecific interactions. Conservation Biology, 28:1342–1348 https://doi.org/10.1111/cobi.12304

Marmor, L., Tõrra, T., Saag, L., & Randlane, T. (2011). Effects of forest continuity and tree age on epiphytic lichen biota in coniferous forests in Estonia. Ecological Indicators, 11:1270–1276 doi:https://doi.org/10.1016/j.ecolind.2011.01.009

Matos, P., Pinho, P., Aragon, G., Martínez, I., Nunes, A., Soares, A. M., Branquinho, C. (2015). Lichen traits responding to aridity. Journal of Ecology, 103:451–458 doi: 10.1111/1365-2745.12364

Matos, P., Geiser, L., Hardman, A., Glavich, D., Pinho, P., Nunes, A., Soares, A.M., & Branquinho, C. (2017). Tracking global change using lichen diversity: towards a global-scale ecological indicator. Methods in Ecology and Evolution, 8: 788–798. https://doi.org/10.1111/2041-210X.12712

McCune, B. (2000). Lichen communities as indicators of forest health. The Bryologist, 103:353–356 https://doi.org/10.1639/0007-2745(2000)103[0353:LCAIOF]2.0.CO;2

Millennium ecosystem assessment M (2005). Ecosystems and human well-being vol 5. Island press Washington, DC,

Miller, J.E.D., Villella, J., Stone, D., & Hardman, A. (2020). Using lichen communities as indicators of forest stand age and conservation value. Forest Ecology and Management, 475:118436 doi:https://doi.org/10.1016/j.foreco.2020.118436

Miller, M. A., Pfeiffer, W., & Schwartz, T. (2010). Creating the CIPRES Science Gateway for inference of large phylogenetic trees. Gateway Computing Environments Workshop, 2010, 1–8. doi: 10.1109/GCE.2010.5676129

Molins, A., Moya, P., Muggia, L., & Barreno, E. (2021). Thallus Growth Stage and Geographic Origin Shape Microalgal Diversity in Ramalina farinacea Lichen Holobionts. Journal of Phycology, 57:975–987 doi:https://doi.org/10.1111/jpy.13140

Moya, P., Molins, A., Martínez-Alberola, F., Muggia, L., & Barreno, E. (2017). Unexpected associated microalgal diversity in the lichen Ramalina farinacea is uncovered by pyrosequencing analyses. PLOS ONE, 12:e0175091 doi:10.1371/journal.pone.0175091

Muggia, L., Nelsen, M. P., Kirika, P. M., Barreno, E., Beck, A., Lindgren, H., Lumbsch, H.T., Leavitt, S.D., (2020). Formally described species woefully underrepresent phylogenetic diversity in the common lichen photobiont genus Trebouxia (Trebouxiophyceae, Chlorophyta): an impetus for developing an integrated taxonomy. Molecular phylogenetics and evolution, 149, 106821. https://doi.org/10.1016/j.ympev.2020.106821

Muggia, L., Pérez-Ortega, S., Kopun, T., Zellnig, G., & Grube, M. (2014). Photobiont selectivity leads to ecological tolerance and evolutionary divergence in a polymorphic complex of lichenized fungi. Annals of Botany, 114:463–475 doi:10.1093/aob/mcu146

Nanda, S. A., Haq, M. U., Singh, S. P., Reshi, Z. A., Rawal, R. S., Kumar, D., Bischt, K., Upadhyay, S., Upreti, D.K., & Pandey, A. (2021). Species richness and β-diversity patterns of macrolichens along elevation gradients across the Himalayan Arc. Scientific Reports, 11(i1), 1–15. https://doi.org/10.1038/s41598-021-99675-1

Nascimbene, J., & Marini, L. (2015). Epiphytic lichen diversity along elevational gradients: biological traits reveal a complex response to water and energy. Journal of Biogeography, 42:1222–1232 doi:10.1111/jbi.12493

Nash, T.H. (1996). Lichen biology. Cambridge University Press,

Nelsen, M.P., & Gargas, A. (2008). Dissociation and horizontal transmission of codispersing lichen symbionts in the genus Lepraria (Lecanorales: Stereocaulaceae). New phytologist, 177:264–275 https://doi.org/10.1111/j.1469-8137.2007.02241.x

Nelson, P.R., McCune, B., Roland, C., & Stehn, S. (2015). Non-parametric methods reveal non-linear functional trait variation of lichens along environmental and fire age gradients. Journal of Vegetation Science, 26:848–865 doi:https://doi.org/10.1111/jvs.12286

Nordén, J., Penttilä, R., Siitonen, J., Tomppo, E., & Ovaskainen, O. (2013). Specialist species of wood-inhabiting fungi struggle while generalists thrive in fragmented boreal forests. Journal of Ecology, 101:701–712 doi:https://doi.org/10.1111/1365-2745.12085

Ollerton, J. (2006). “Biological barter”: patterns of specialization compared across different mutualisms Plant-pollinator interactions: from specialization to generalization University of Chicago Press, Chicago:411–435

Otálora, M.A.G., Salvador, C., Martínez, I., & Aragón, G. (2013). Does the Reproductive Strategy Affect the Transmission and Genetic Diversity of Bionts in Cyanolichens? A Case Study Using Two Closely Related Species. Microbial Ecology, 65:517–530 doi:10.1007/s00248-012-0136-5

Ott, S. (1987). The juvenile development of lichen thalli from vegetative diaspores. Symbiosis, 3:57–74

Paoli, L., Pinho, P., Branquinho, C., Loppi, S., & Munzi, S. (2017). The influence of growth form and substrate on lichen ecophysiological responses along an aridity gradient. Environmental Science and Pollution Research, 24:26206–26212 doi: https://doi.org/10.1007/s11356-017-9361-2

Peksa, O., Škaloud, P. (2011). Do photobionts influence the ecology of lichens? A case study of environmental preferences in symbiotic green alga Asterochloris (Trebouxiophyceae). Molecular Ecology, 20: 3936–3948. doi: https://doi.org/10.1111/j.1365-294X.2011.05168.x

Pérez-Ortega, S., Garrido-Benavent, I., Grube, M., Olmo, R., & de los Ríos, A. (2016). Hidden diversity of marine borderline lichens and a new order of fungi: Collemopsidiales (Dothideomyceta). Fungal Diversity, 80(1), 285–300. https://doi.org/10.1007/s13225-016-0361-1

Pérez-Ortega, S., Ortiz-Álvarez, R., Allan Green, T.G., & de los Ríos, A. (2012). Lichen myco- and photobiont diversity and their relationships at the edge of life (McMurdo Dry Valleys, Antarctica). FEMS Microbiology Ecology, 82:429–448 doi:10.1111/j.1574-6941.2012.01422.x

Pinho, P., Bergamini, A., Carvalho, P., Branquinho, C., Stofer, S., Scheidegger, C., Máguas, C. (2012). Lichen functional groups as ecological indicators of the effects of land-use in Mediterranean ecosystems. Ecological Indicators, 15:36–42. DOI: https://doi.org/10.1016/j.ecolind.2011.09.022.

Plummer, M., Best, N., Cowles, K., & Vines, K. (2006). CODA: convergence diagnosis and output analysis for MCMC. R News, 6(1) pp. 7–11.

Poisot, T., Canard, E., Mouillot, D., Mouquet, N., & Gravel, D. (2012). The dissimilarity of species interaction networks. Ecology Letters, 15:1353–1361. https://doi.org/10.1111/ele.12002

Poisot, T., Stouffer, D.B., & Gravel, D. (2015). Beyond species: why ecological interaction networks vary through space and time. Oikos, 124:243–251. doi:https://doi.org/10.1111/oik.01719

Rambaut, A. (2012). FigTree v1. 4. Molecular evolution, phylogenetics and epidemiology. Edinburgh: University of Edinburgh, Institute of Evolutionary Biology. Available at: http://tree.bio.ed.ac.uk/software/figtree/

Rambaut, A., Drummond, A. J., Xie, D., Baele, G., & Suchard, M. A. (2018). Posterior summarization in Bayesian phylogenetics using Tracer 1.7. Systematic biology, 67(5), 901–904. https://doi.org/10.1093/sysbio/syy032

Reif, J., Horák, D., Krištín, A., Kopsová, L., & Devictor, V. (2016). Linking habitat specialization with species’ traits in European birds. Oikos, 125:405–413 doi:https://doi.org/10.1111/oik.02276

Rivas Plata, E., Lücking, R., & Lumbsch, H.T. (2008). When family matters: an analysis of Thelotremataceae (Lichenized Ascomycota: Ostropales) as bioindicators of ecological continuity in tropical forests. Biodiversity and Conservation, 17:1319–1351 doi:10.1007/s10531-007-9289-9

Rolshausen, G., Dal Grande, F., Sadowska-Des, A.D., Otte, J., & Schmitt, I. (2018). Quantifying the climatic niche of symbiont partners in a lichen symbiosis indicates mutualist-mediated niche expansions. Ecography, 41:1380–1392 doi:https://doi.org/10.1111/ecog.03457

Rosenzweig, M.L. (1995). Species diversity in space and time. Cambridge University Press,

Sanders, W.B. (2005). Observing microscopic phases of lichen life cycles on transparent substrata placed in situ. The Lichenologist, 37:373–382 doi:10.1017/S0024282905015070

Sanders, W.B., & Lücking, R. (2002). Reproductive strategies, relichenization and thallus development observed in situ in leaf-dwelling lichen communities. New Phytologist, 155:425–435 doi:https://doi.org/10.1046/j.1469-8137.2002.00472.x

Santamaría, L., & Rodríguez-Gironés, M.A. (2007). Linkage Rules for Plant–Pollinator Networks: Trait Complementarity or Exploitation Barriers?. PLOS Biology, 5:e31 doi:10.1371/journal.pbio.0050031

Schoch, C. L., Seifert, K. A., Huhndorf, S., Robert, V., Spouge, J. L., Levesque, C. A., & Chen, W., (2012). Nuclear ribosomal internal transcribed spacer (ITS) region as a universal DNA barcode marker for Fungi. Proceedings of the National Academy of Sciences, 109(16), 6241–6246. https://doi.org/10.1073/pnas.1117018109

Singh, G., Dal Grande, F., Divakar, P. K., Otte, J., Leavitt, S. D., Szczepanska, K., Crespo, A., Rico, V.J., Aptroot, A., Cáceres, M.E.dS., Lumbscht H.T., & Schmitt, I. (2015). Coalescent-based species delimitation approach uncovers high cryptic diversity in the cosmopolitan lichen-forming fungal genus Protoparmelia (Lecanorales, Ascomycota). PLoS One, 10(5). https://doi.org/10.1371/journal.pone.0124625

Solé, R.V., & Montoya, M. (2001). Complexity and fragility in ecological networks. Proceedings of the Royal Society of London Series B: Biological Sciences 268:2039–2045 doi:doi:10.1098/rspb.2001.1767

Svoboda, D., Peksa, O., & Veselá, J. (2010). Epiphytic lichen diversity in central European oak forests: Assessment of the effects of natural environmental factors and human influences. Environmental Pollution, 158:812–819 doi:https://doi.org/10.1016/j.envpol.2009.10.001

Thompson, J.N. (1988). Variation in interspecific interactions. Annual review of ecology and systematics, 19:65–87

Thüs, H., Muggia, L., Pérez-Ortega, S., Favero-Longo, S. E., Joneson, S., O’Brien, H., Nelsen, P.M., Duque-Thüs, R., Grube, M., Friedl, T., Brodie, J., Andrew, C.J., Lücking, R., Lutzoni, F., & Gueidan, C. (2011). Revisiting photobiont diversity in the lichen family Verrucariaceae (Ascomycota). European Journal of Phycology, 46(4), 399–415. https://doi.org/10.1080/09670262.2011.629788

Toju, H., Tanabe, A.S., Yamamoto, S., & Sato, H. (2012). High-Coverage ITS Primers for the DNA-Based Identification of Ascomycetes and Basidiomycetes in Environmental Samples. PLOS ONE, 7:e40863 doi:10.1371/journal.pone.0040863

Tretiach, M., Crisafulli, P., Pittao, E., Rinino, S., Roccotiello, E., & Modenesi, P. (2005). Isidia ontogeny and its effect on the CO2 gas exchanges of the epiphytic lichen Pseudevernia furfuracea (L.) Zopf. The Lichenologist, 37:445–462 doi:10.1017/S0024282905014982

Tripp, E.A., & Lendemer, J.C. (2018). Twenty-seven modes of reproduction in the obligate lichen symbiosis. Brittonia, 70:1–14 doi:10.1007/s12228-017-9500-6

Trøjelsgaard, K., Jordano, P., Carstensen, D.W., & Olesen, J.M. (2015). Geographical variation in mutualistic networks: similarity, turnover and partner fidelity. Proceedings of the Royal Society B: Biological Sciences, 282:20142925 doi:doi:10.1098/rspb.2014.2925

Tschermak-Woess, E. (1988). The algal partner. In CRC handbook of lichenology 1:39–92

Tylianakis, J.M., Didham, R.K., Bascompte, J., & Wardle, D.A. (2008). Global change and species interactions in terrestrial ecosystems. Ecology letters, 11:1351–1363 https://doi.org/10.1111/j.1461-0248.2008.01250.x

Tylianakis, J.M., & Morris, R.J. (2017). Ecological Networks Across Environmental Gradients. Annual Review of Ecology, Evolution, and Systematics, 48:25–48 doi:10.1146/annurev-ecolsys-110316-022821

Valiente-Banuet, A., Aizen, M.A., Alcántara, J.M., Arroyo, J., Cocucci, A., Galetti, M., García, M.B., García, D., Gómez, J.M., Jordano, P., Medel, R., Navarro, L., Obeso, J.R., Oviedo, R., Ramírez, N., Rey, P.J., Traveset, A., Verdú, M., & Zamora, R. (2015). Beyond species loss: the extinction of ecological interactions in a changing world. Functional Ecology, 29: 299–307. https://doi.org/10.1111/1365-2435.12356

Walser, J-C. (2004). Molecular evidence for limited dispersal of vegetative propagules in the epiphytic lichen Lobaria pulmonaria. American Journal of Botany, 91:1273–1276 doi:https://doi.org/10.3732/ajb.91.8.1273

Webb, C.T., Hoeting, J.A., Ames, G.M., Pyne, M.I., & LeRoy Poff, N. (2010). A structured and dynamic framework to advance traits-based theory and prediction in ecology. Ecology Letters, 13:267–283 doi:https://doi.org/10.1111/j.1461-0248.2010.01444.x

Wickham, H. (2016). ggplot2: elegant graphics for data analysis. Springer,

Wilcove, D.S., McLellan, C.H., & Dobson, A.P. (1986). Habitat fragmentation in the temperate zone. Conservation biology, 6:237–256

Wornik, S., & Grube, M. (2010). Joint Dispersal Does Not Imply Maintenance of Partnerships in Lichen Symbioses. Microbial Ecology, 59:150–157 doi:10.1007/s00248-009-9584-y

Yahr, R., Vilgalys, R., & DePriest, P.T. (2006). Geographic variation in algal partners of Cladonia subtenuis (Cladoniaceae) highlights the dynamic nature of a lichen symbiosis. New phytologist, 171:847–860 https://doi.org/10.1111/j.1469-8137.2006.01792.x

